# Discordant population structure inferred from male- and female-type mtDNAs from *Macoma balthica*, a bivalve species characterized by doubly uniparental inheritance of mitochondria

**DOI:** 10.1101/2022.02.28.479517

**Authors:** Sabrina Le Cam, Julie Brémaud, Vanessa Becquet, Valérie Huet, Emmanuel Dubillot, Pascale Garcia, Amélia Viricel, Sophie Breton, Eric Pante

## Abstract

Doubly Uniparental Inheritance (DUI) of mitochondria is a remarkable exception to the Strictly Maternal Inheritance (SMI) in metazoans. In species characterized by DUI --almost exclusively gonochoric bivalve mollusks--females (F) transmit mitochondria to offspring of both sexes, while males (M) pass on their mitochondria exclusively to their sons. Under DUI, males are heteroplasmic, somatic tissues containing F-transmitted mtDNA and gametic cells containing M-transmitted mtDNAs. The aforementioned transmission routes make M- and F-transmitted mtDNA interesting as sex-specific markers which can differ in their effective population sizes, mutation rates, and selective constraints. For these reasons, looking at both markers can provide significant insights into the genetic structure of populations and investigate its determinants. In this study, we document differences in genetic diversity, divergence, inter-populational differentiation and biogeographic structure between M- and F-type *cox1* mt genes in the Baltic tellin (*Macoma balthica*) to test whether *cox1m* and *cox1f* genes bear the marks of similar phylogeographic histories. These markers were sequenced for 302 male individuals sampled from the North Sea to the Gironde Estuary (Southern France) encompassing the intra-subspecific *M. b. rubra* hybrid zone in the Gulf of Saint Malo. Both genes supported a scenario of cladogenesis of *M.b. rubra* clades prior to the last glacial maximum. Nucleotide diversity and net divergence were over twice higher in *cox1m* compared to *cox1f*. Genetic differentiation between northern and southern populations was nearly 3 times higher at *cox1m* compared to *cox1*f (global Φ_ST_ = 0.345 and 0.126 respectively) and the geographic localization of the strongest genetic break significantly differed between the markers (Finistère Peninsula at *cox1f*; Cotentin Peninsula at *cox1m*, ∼250 km apart). A higher mutation rate, relaxed negative selection and differences in effective population sizes (depending on locations) at *cox1m* could explain differences in population genetic structure. As both F- and M-type mtDNAs interact with nuclear genes for oxidative phosphorylation and ATP production, geographical discordances in genetic clines in a context of secondary contact could be linked to mito-nuclear genetic incompatibilities.

## Introduction

Some species show a remarkable exception to the maternal inheritance of mitochondria in metazoans: the doubly uniparental mode of inheritance (DUI). In this system, both males and females can transmit their mitochondria. The former transmits female-inherited F-type mitochondria to all their progeny and the latter pass on “male-inherited” (M-type) mitochondria to their male offspring, where the male mitogenomes (mt) are quartered in male germ line and gametes (reviewed in Zouros, 2013). To date, DUI has only been discovered in the class Bivalvia, with over 100 DUI species (Gusman et al., 2016) among the *c.a.* 11,000 (Huber, 2010; Huber et al., 2015). They are all gonochoric, except for the hermaphroditic mussel *Semimytilus algosus* (Lubośny et al., 2020). More than a simple peculiarity, DUI is suspected to play a role in sex-determination and gonad differentiation (Zouros, 2000; Breton et al., 2011; Guerra et al., 2017; Capt et al., 2018, 2019) and could well be involved in population structure through intrinsic (*e.g.* genetic incompatibilities; Saavedra et al., 1996) and extrinsic (*e.g.* selection and demography, Stewart et al., 1996) factors.

In DUI species, divergence between F-type and M-type mitogenomes is variable but generally high, ranging from 6 to over 50% (reviewed in Breton et al., 2007; Gusman et al., 2016), which questions the maintenance of mito-nuclear genetic coadaptation. Indeed, both F- and M-type mitochondria can be found in males and females but in majority, females are homoplasmic for the F-type mtDNA whereas males are heteroplasmic, accommodating two highly divergent mitogenomes (F-type in somatic tissues and M-type in sperm). The presence of the M-type mtDNA in somatic tissues is considered as a paternal leak due to elimination or segregation failure of sperm mitochondria in female or male embryos, respectively (Milani et al., 2012). Both F- and M-type mt lineages show rapid molecular evolution compared to other animals, the M-type mtDNA usually evolving faster than the F-type mtDNA (Zouros, 2013). Coevolution and coadaptation of mitochondrial and nuclear genes are required for efficient cellular energy production (*i.e.* oxidative phosphorylation OXPHOS) and mito-nuclear genetic incompatibilities (MNIs) can lead to a desynchronization of this machinery (Burton & Barreto, 2012; Burton et al., 2013). DUI could, therefore, bear on the maintenance of genetic structure among populations of highly dispersive bivalve species at small spatial scales and provide key insight into the establishment and maintenance of local adaptation.

Indeed, barriers to gene flow can arise and be maintained by a multitude of environmental and/or intrinsic factors (Barberousse et al., 2010), from ecological isolation to genetic incompatibilities. Hybrid zones, which correspond to regions spatially separating genetic stocks, are “natural laboratories” to study the interactions between intrinsic barriers and the environment, and the processes of adaptation and speciation.

*Macoma balthica*, a species in which DUI has recently been detected (Pante et al., 2017; Capt et al., 2020), is a noteworthy model species to study hybrid zones in marine ecosystems (Riginos & Cunningham, 2007; Strelkov et al., 2007). It has a wide distribution range spanning from the west pacific coasts, in Japan and from Alaska to Oregon (USA, Luttikhuisen et al., 2003) to the North Atlantic, where the species is found in the west from Arctic to Newfoundland (Layton et al., 2016) and in the east from the north of Russia (Hummel et al., 1997) to the Arcachon Basin (Hily, 2013 and this publication). The succession of glaciation and inter-glaciation periods has resulted in colonization of the Atlantic marked by repeated isolation, invasion, and re-colonization events (Nikula et al., 2007). These episodic colonization events have created multiple opportunities for secondary contacts between different genetic stocks and the establishment of several hybrid zones in the North Atlantic, where two subspecies of *M. balthica* co-occur. A Pacific lineage (*M. b. balthica*) is present in the Baltic Sea and the White Sea, and an Atlantic lineage (*M. b. rubra*) is found in the Norwegian Sea, the North Sea and along the British coasts, down to the southern range limit of the species (Luttikhuizen et al., 2003; Nikula et al., 2008; Väinölä, 2003). In Europe, two genetic breaks were described: an inter-subspecific break occurs between *M. b. balthica* and *M. b. rubra* in the transition zone from the North Sea to the Baltic Sea (Luttikhuizen et al., 2012; Nikula et al., 2007, 2008), and an intra-subspecific break within *M. b. rubra* in the Gulf of Saint-Malo (Brittany, France). In the latter case, southern populations of *M. b. rubra* (distributed between the southern range limit and the aforementioned break) exhibit F-type mtDNA signatures consistent with long-term isolation in the glacial refugium of the Bay of Biscay: high genetic diversity relative to previously glaciated areas, high prevalence of private alleles, and sharp genetic differentiation from northern populations (Becquet et al., 2012). Deciphering signal of primary intergradation from secondary contact in hybrid zones is challenging; we have therefore gathered sequences from both F and M mitogenomes to estimate the timing of cladogenesis in the *Macoma* species complex, particularly in the *M. b. rubra* hybrid zone. Clarifying the phylogeographic context will allow us to focus on the effects of DUI on the evolution of *M. b. rubra* lineages. Indeed, multiple nuclear genes involved in the oxidative phosphorylation (OXPHOS) system of mitochondria were detected as significantly differentiated among southern and northern populations (Pante et al., 2012, 2019); DUI could therefore play a role in the maintenance of genetic barriers in *M. b. rubra*.

## Materials and Methods

### Sampling

Individuals were collected from a total of 14 sampling sites ranging from Arcachon (southern range limit of the species, France) to Le Crotoy (Somme Bay, northern France) and from Kruiningen (the Netherlands) to Sylt (Germany) (Table S1). Individuals from sampling sites on the French coasts were treated as follows: 70 to 100 adults from 11 mm to 23 mm were randomly collected live at sexual maturity between 4th and 23rd of April 2018 at 10 locations ranging from the Bay of Biscay in southern France to Somme Bay (Table S1). Individuals were then held in aquaria until dissection at the LIENSs laboratory (LIttoral ENvironnement et Sociétés) in La Rochelle, France, with water temperature maintained at 10°C. They were fed with a multispecific microalgal mixture every other day and dead or dying individuals were removed daily. For each individual, the adductor muscle was carefully severed to separate the two valves, without damaging the gonad, and a sample of the mantle was taken. Sex and gonadal maturation stage were then determined with a dissecting microscope (x100 to x400). Two types of tissue samples were collected: a gonadic sample and a somatic sample (mantle). All tissue samples were flash-frozen in liquid nitrogen and then stored at -80°C until DNA extraction. At each step of this protocol, dissecting tools and bench surfaces were thoroughly cleaned in successive baths of 10 % bleach, demineralized water, and 95 % EtOH as to avoid DNA cross-contamination. Using this dataset, we checked that male mitochondria were limited to male gonads (*i.e.* absent from other male tissues and from females); thus making molecular sexing permissible (Le Cam et al., 2023).

For the other sampling sites, whole individuals had been collected for a previous study (Becquet et al., 2012 and the BIOCOMBE project) and were preserved in 95% EtOH. A 3 mm^3^ piece of tissue was sampled from the gonad, gDNA was purified and quantified as detailed below, and PCR amplifications were attempted with both the *cox1m* and *cox1f* primers. Individuals for which both gene regions could be amplified were considered as males and included in subsequent analyses. For phylogenetic inferences, additional samples were sequenced to grasp as much diversity as possible within the *Macoma* species complex, from the NW, NE Atlantic and the NE Pacific (table S1). They were combined to available data from Nikula et al., (2007), Becquet et al., (2012), Layton et al., (2016) and Yurchenko et al., (2018)

### Molecular Analyses

Total DNA was extracted with the Nucleospin Tissue Kit (Macherey Nagel), following the manufacturer’s instructions. In Saint-Vaast-la-Hougue (VAA) and Mont-Saint-Michel (MSM), most specimens were sexually undifferentiated at the time of sampling. Therefore, for these samples and for the samples from all the other “non-French” localities, sex was determined by checking for the amplification of both male and female *cox1* markers in gonad DNA. DNA purity and potential contaminants were checked using a NanoDrop2000.

Specific primers were designed to independently amplify the 5’ portion of the *cox1* gene of the M-type and F-type mitogenomes (*cox1m* and *cox1f* respectively). To do that, both mitogenomes published in Capt et al., 2020 were used and they came from *M. b. rubra* specimens sampled in Aytré, France (clade b1b). We also used *cox1* sequences retrieved from transcriptomic data (Pante et al., 2012) from specimens sampled in Aytré, Somme Bay (France, b1a clade) and Gdansk (Poland, b2/b3 clade). Polymerase chain reaction (PCR) amplifications of *cox1f* and *cox1m* mitochondrial DNA regions were performed using the following primers (*cox1m*: cox1m14641F ATAGCTGGCCTGGTGTTTAGG, cox1m15560R TTGGACCCTTTCGAGCCAAG; *cox1f*: cox1f5434F TTAGTGACTTCACACGGTTTGC, cox1f6032R TGGGAAATAATCCCAAACCCG). Amplifications were realized in a total volume of 25 µL, with 0.1 µL of Taq Polymerase 5 U.µL^-1^ (TaKaRa Ex Taq® Kit MgCl_2_ Free Buffer) (TaKaRa Ex Taq, TaKaRa Bio, Shiga, Japan), 2.5 µL PCR Buffer 10X, 1.5 µL MgCl_2_ 25 mM, 1 µL dNTP 2.5 mM each, 0.6 µL each primer 10 µM and 17.7 Milli-Q water, and 1 to 20 ng of template DNA. SensoQuest thermal cyclers (Göttingen, Germany) were used to perform the following PCR cycling profiles: (i) for the *cox1f*, 2 min of initial denaturation at 94°C, then 30 cycles consisting of 45 sec at 94°C followed by 30 sec of annealing at 57°C and 40 sec of elongation at 72°C, and a 5 min final step of elongation at 72°C, and (ii) for the *cox1m*, 2 min at 94°C, then 30 cycles of 30 sec at 94°C, 30 sec at 60°C and 55 sec at 72°C, and a final elongation for 5 min at 72°C. PCR success (*i.e.* amplicon concentration, specificity and absence of amplicons for PCR and extraction negative controls), was tested on 1% agarose gels. PCR products were then sent to Eurofins GATC Biotech GmbH (Konstanz, Germany) to be purified and Sanger sequencing was performed in both directions.

### Sequence Analyses

All sequences of *cox1f* sequences and *cox1m* fragments were quality-controlled and aligned in Geneious Prime 10.2.6 (Kearse et al., 2012). Clade names follow Nikula et al., (2007). Consensus sequences were trimmed to 479 bp and 676 bp for *cox1f* and *cox1m*, respectively. *cox1m* sequences, trimmed at the same length as *cox1f* (479bp), were used for further comparative analyses. Haplotype sequences and their frequencies were computed in R v3.3.3 (R Core Team, 2017) using the *pegas* package v0.11.(Paradis, 2010). In order to model intraspecific relationships among haplotypes, for both *cox1f* and *cox1m*, median-joining networks (Bandelt et al., 1999) were constructed with PopART v1.7 software (Leigh & Bryant, 2015). For each marker, we constructed a maximum-likelihood phylogenetic tree using IQ-TREE software (Nguyen et al., 2015) with 1,000 bootstrap replicates, and nucleotide acid-substitution models were inferred using ModelFinder with Bayesian information criteria (Nguyen et al., 2015; Kalyaanamoorthy et al., 2017). For *cox1f*, data from Nikula et al., (2007), Becquet et al., (2012), Layton et al., (2016) and Yurchenko et al., (2018) were used for phylogenetic inference. For *cox1m*, we also sequenced data from the *Macoma* species complex, that is *M. petalum* (clade A) and *M balthica balthica (*clades C and D). From these genealogical relationships, mitochondrial lineages were defined (haplogroups) and their population relative frequency estimated.

We used the net distance approach to estimate the timing of cladogenesis events within the *Macoma* tree.We used the same dataset as described for phylogenetic reconstruction, estimated the most likely model of nucleotide substitution in IQTREE. In MEGA 11 (Tamura et al., 2021), we calculated net distances using the Tamura-Nei model (Tamura & Nei, 1993) as it was the closest model identified by IQTREE. The rate variation among sites was modeled with a gamma distribution (shape parameter = 0.17). The differences in the composition bias among sequences were considered in evolutionary comparisons (Tamura & Kumar, 2002). All codon positions were included, using the pairwise deletion option for missing data. We used 115, 479 bp-long sequences for *cox1f* and 102, 676 bp-long sequences for *cox1m*. The substitution rates for *cox1f* and *cox1m* were then estimated using the net distance described above and the calibration dates proposed in Nikula et al., (2007) for the trans-Atlantic divergence between rubra and balthica. 3.5My corresponds to the opening of the Bering Strait, the earliest time of invasion of *Macoma* into the Atlantic. 2My corresponds to the earliest known fossils of *Macoma balthica* in Europe (Norton & Spaink, 1973). This strategy was also used by Wares & Cunningham, (2001) and Riginos & Cunningham, (2007), and Riginos et al., (2004) in the context of DUI.

Genetic diversity indices and population differentiation analyses at *cox1f* and *cox1m* were estimated only for sampling sites with a minimum sample size of 10 individuals. Genetic diversity indices were calculated using Arlequin v3.5.2.2 (Excoffier et al., 1992): haplotype number (H), haplotype diversity (Hd) and nucleotide diversity (π Tajima, 1983). The differences between the distributions of each genetic diversity statistic (π, Hd) were statistically evaluated using Wilcoxon rank tests for paired data. Genetic differentiation was estimated in Arlequin using the *Φ*_ST_ fixation index and its statistical significance was tested by performing 10,000 permutations (Excoffier et al., 1992).We performed maximum-likelihood geographical cline analyses with the r package HZAR 0.2-9 (Derryberry et al., 2014) together with cline centers and cline widths estimates. Clines were fitted on haplogroup frequencies and geographical distances from the most northern sample sites (SYL). For each haplogroup frequencies, two models were fitted and compared to the null model (no cline): I (fixed scaling based on minimum and maximum observed frequencies and no exponential tails) and II (free scaling and no exponential tails). We performed model selection using AICc scores to determine the preferred model and then extracted the ML model parameters. Geographical distances along the coast were computed using the marmap 1.0.10 R package (Pante & Simon-Bouhet, 2013).

An analysis of molecular variance (AMOVA) was carried out to evaluate the genetic structure among geographic regions and overall sampling sites using 10,000 permutations with Arlequin v3.5.2.2 (Excoffier et al., 1992). We used jModeltest2 (Guindon & Gascuel, 2003; Darriba et al., 2012) to choose the Tamura-Nei model of nucleotide substitution (with gamma shape parameters of 0.1150 and 0.1450 for *cox1f* and *cox1m*, respectively) as a measure of haplotype distances in Arlequin v3.5.2.2. Based on discordances in the distribution of the genetic diversity at *cox1f* and *cox1m*, several hierarchical groupings of subpopulations were tested for each marker to determine the best geographical delineation of genetic structure (Table 2: 3 groups (CRI,KRU / CRO,VAA,MSM,BRI / MAH,BRE,AYT,FOU,ARC); 2 groups A (CRI,KRU,CRO,VAA / MSM,BRI,MAH,BRE,AYT,FOU,ARC); 2 groups B (CRI,KRU,CRO,VAA,MSM,BRI / MAH,BRE,AYT,FOU,ARC)). Different measures of populations differentiation (pairwise and AMOVA) were also carried out using conventional F-statistics.

For the *cox1f* and *cox1m* markers, mismatch distribution analyses were performed to infer deviations from long-term population stability. We considered two genetically-homogeneous groups (as in Marko et al., 2010): a “North” group composed of individuals from KRU, SYL, WIL and CRI, and a “South” group with individuals from AYT, ARC, BRE and FOU. As detailed in the Results, no genetic sub-structure was detected within these two groups using pairwise *Φ*_ST_. Observed distributions were compared to distributions expected under constant size and growth/decline models. Raggedness r (Harpending, 1994) and Ramos-Onsins and Rozas’ R_2_ (Ramos-Onsins & Rozas, 2002) statistics were estimated, and their significance along with confidence intervals were computed using coalescent simulations (10 000 replicates). These analyses were carried out with DNAsp6 (Rozas et al., 2017). To test for differences in selective pressure on the female and male mitochondrial sequence sets, Fu’s Fs (Fu, 1997) and Tajima’s D (Tajima, 1989) test for deviations from the neutral theory model for a population of constant size was calculated in Arlequin v3.5.2.2 (Excoffier et al., 1992) for each marker and group. Their statistical significance was tested with 10,000 permutations. Finally, relative differences in effective population size between both markers were estimated with the theta ratio (Watterson, 1975) among genetically homogenous groups of samples (as described above for mismatch distributions). Theta indices, based on the number of segregating sites were estimated using the *theta.s* function of the pegas 1.3 R package (Paradis, 2010).

## Results

While this study focuses on the differences in genetic diversity and population structure at the *cox1f* and *cox1m* markers, from the North Sea to the southern limit of the distribution range, we first investigated the overall phylogeographic context of these lineages. Phylogenetic inferences from the *cox1f* dataset combined with Nikula et al., (2007), Becquet et al., (2012), Layton et al., (2016) and Yurchenko et al., (2018) data revealed that our English Channel / Bay of Biscay dataset is only composed of the *M. balthica rubra* lineage (Supplementary material figure S1, locations of the samples are presented in figure S3). The phylogenetic tree and the haplotype network were congruent: the “northern” haplogroup corresponded to the b2/b3 *M. b. rubra* lineage and our data revealed 2 sub-lineages within the b1 lineage, b1a and b1b. The phylogeny of *cox1m* revealed a similar pattern, with three clades m1, m2, m3 (Supplementary material Figure S2) and a fourth clade m4 encompassing mainly individuals from the Baltic Sea and originating at the base of the *M. b. rubra* branch. Based on net divergence, we were able to compare the timing of cladogenesis of *cox1m* and *cox1f* clades (Table S2). In particular, the divergence of *cox1m* and *cox1f* clades encountered in the *M. b. rubra* hybrid zone all seem to have occurred before the last glacial maximum (*cox1f* clades b1a and b1b between 106 and 185 ky, *cox1m* clades m1 vs m2/m3 between 30 and 50 ky).

Sexing with optical microscopy on 655 individuals from 14 sampling sites of the French Atlantic coast resulted in the detection of 248 males, 185 females and 222 undifferentiated individuals. Undifferentiated specimens might have released their gametes or resorbed them, as lipid droplets were observed in some of the undifferentiated gonads. Molecular sexing was used to detect males in undifferentiated individuals. Both *cox1m* and *cox1f* were successfully sequenced for 302 male individuals from 14 sampling sites (Table 1). Sampling size per site ranged from 3 to 33 overall sampling sites and from 28 to 33 when only considering sampling sites in France, apart from Arcachon (ARC, N=10). Among the 302 studied male individuals, 34 and 61 haplotypes were found for *cox1f* and *cox1m,* respectively. Among these, 59% were singletons at *cox1f* and 80% at *cox1m*.

**Table 1.**
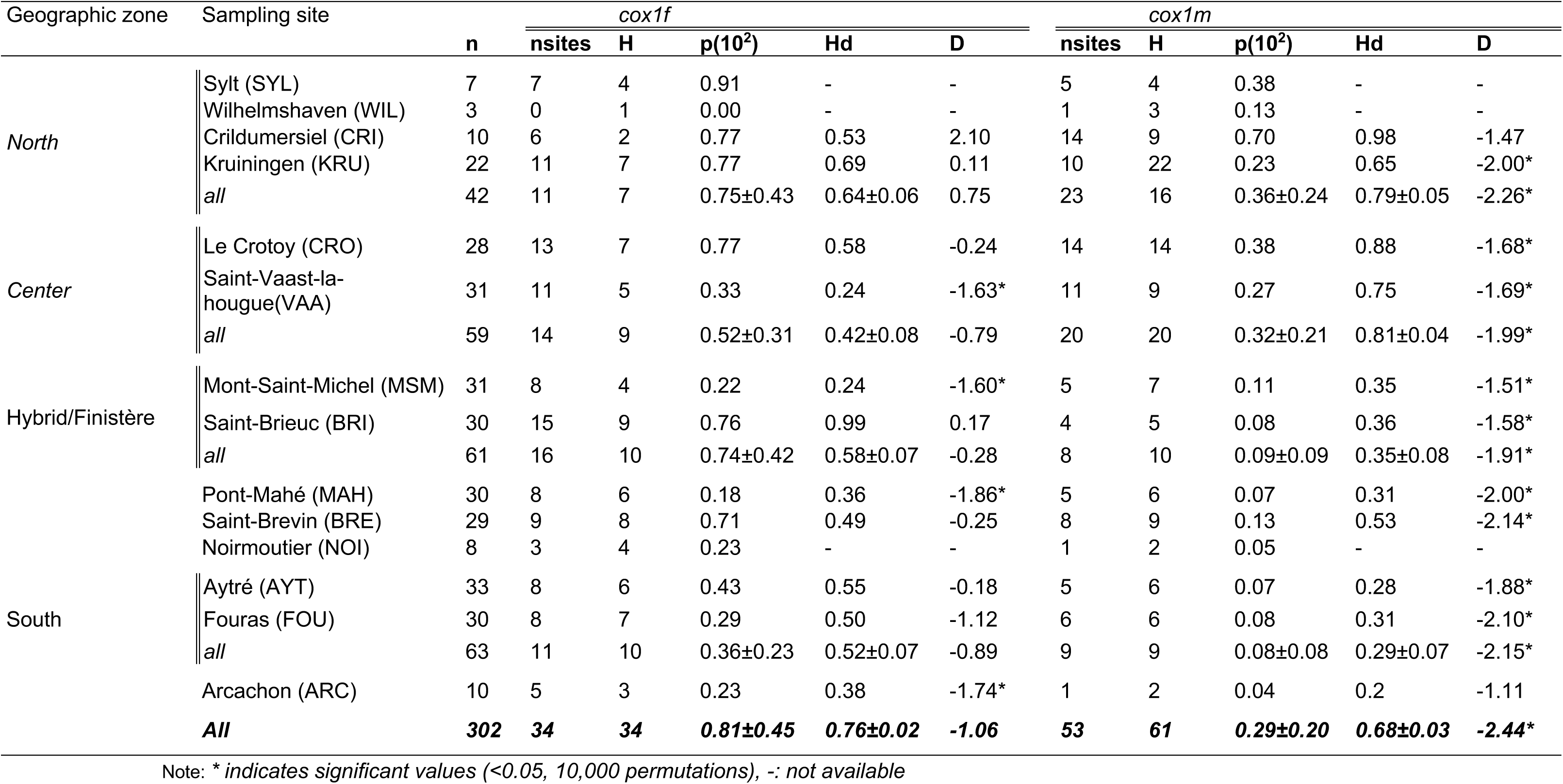
Genetic diversity indices at *cox1f* (479bp) and *cox1m* (479bp) loci for each sampling site: n: number of individuals, nsites: number of polymorphic sites, H: number of haplotypes, p: nucleotide diversity, Hd: Haplotype diversity and D: Tajima’s D coefficient.

Median joining networks showed a similar geographical pattern with three major haplogroups encompassing mainly the northern/central and southern samples (Figure 1A and C). Nonetheless, the networks revealed important differences between markers. First, *cox1m* presented a higher diversity, but a similar divergence level: a maximum of 25 and 26 mutation steps separated the most divergent haplotypes for *cox1f* and *cox1m,* respectively. We estimated maximum divergence rates of 0.027 s/s and 0.028 s/s for the m and f datasets, respectively.

**Figure 1.**
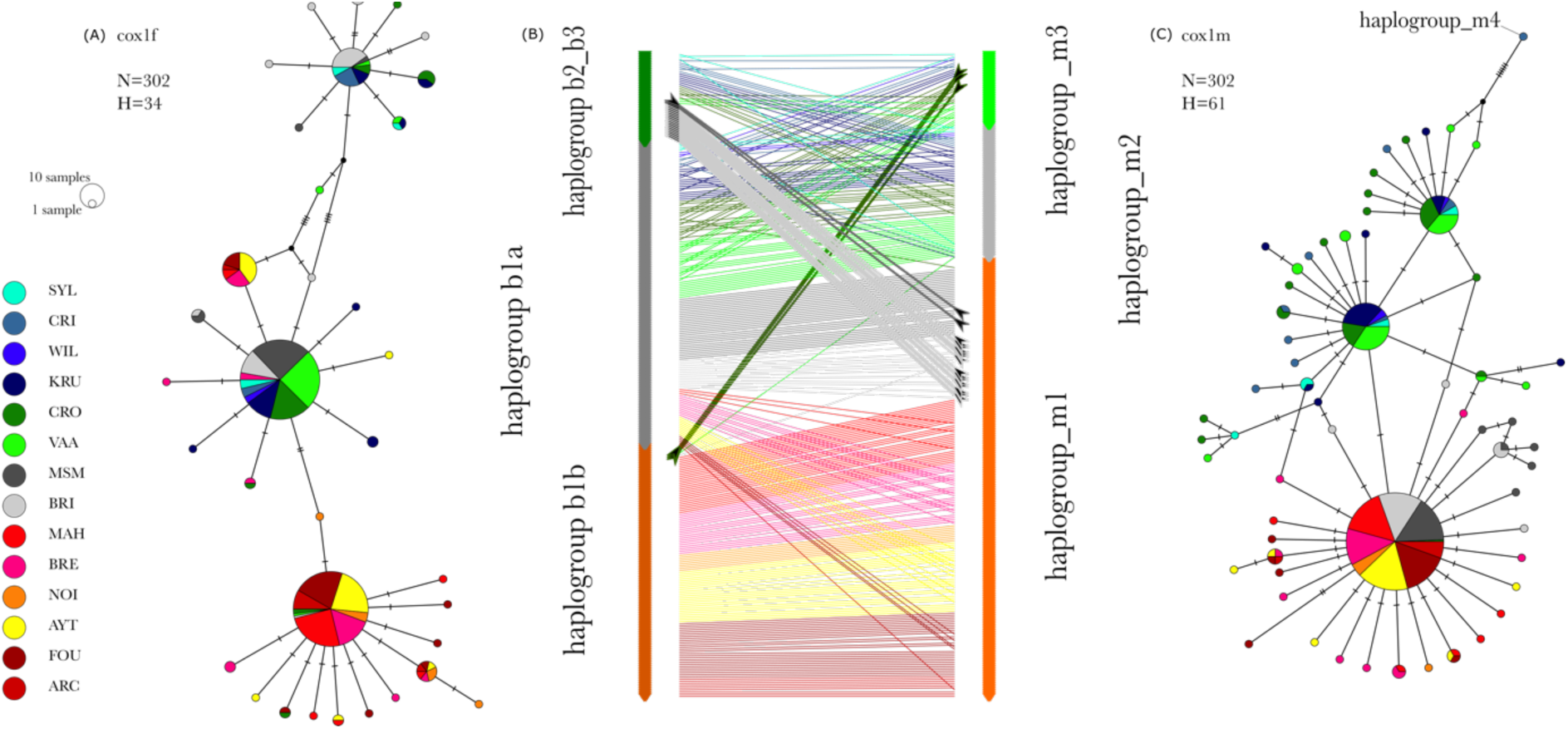
Median joining networks constructed from the haplotypes found among the 302 individuals distributed in 14 sampling sites for (A) the *cox1f* gene and (C) the *cox1m* gene. Haplotype associations among haplogroups are represented with a tanglegram (B) Haplotypes are sorted by haplogroup for each marker and a line represent the connection of the *cox1m* and *cox1f* haplotypes within one individual. The color of the line represents the population where it was sampled and bold lines and arrows highlighting individuals sharing distant haplogroups

The haplogroup relative frequencies showed different spatial patterns (Figure 2A). The *cox1m* haplogroup m3 was present only in the North Sea and east Channel samples while the *cox1f* haplogroup b2/b3 encompassed individuals from the North Sea to the west English Channel (hybrid zone). Also, the haplogroup b1a of *cox1f* was shared among the North Sea, the English Channel and Atlantic samples, whereas the central haplogroup m2 of *cox1m* was found only in the North Sea and the Channel Sea. Interestingly, clear genetic breaks were revealed for each marker, but they were not located at the same geographical location. For *cox1m*, the Cotentin peninsula represented a sharp transition zone between the haplogroups m3/m2 and m1, which was confirmed by pairwise *Φ*_ST_ values (Table S3). Two homogenous populations were observed north of the Cotentin peninsula and south of the Finistère (within group pairwise *Φ*_ST_ ranged from -0.05 to 0.04) and these populations were highly differentiated (between group pairwise *Φ*_ST_ ranged from 0.38 to 0.61). In between, the BRI and MSM sampling sites were very differentiated (between group pairwise *Φ*_ST_ ranged from 0.50 to 0.61) from the North of the Cotentin but they were also weakly yet significantly different from some sites at the South of the Finistère (between group pairwise *Φ*_ST_ ranged from 0.02 to 0.04). For *cox1f*, the pattern was more complex. The haplogroup b1a was composed of haplotypes found in majority at the south of the Finistère (Figure 2) and samples from either side of the Finistère were highly divergent (pairwise *Φ*_ST_ ranging from 0.41 to 0.79, Table S4). On both sides of the Finistère peninsula, significant population substructure was detectable: the Pont-Mahé Bay (MAH) sampling site was mildly differentiated from Saint-Brevin-les-Pins (BRE) and Aytré (AYT) (pairwise *Φ*_ST_ of 0.13 and 0.08 respectively, Table S4). Also, the relative frequencies of the 3 haplogroups were variable in the north of the Finistère peninsula, resulting in a heterogeneous distribution of the genetic diversity (Figure 2).

**Figure 2.**
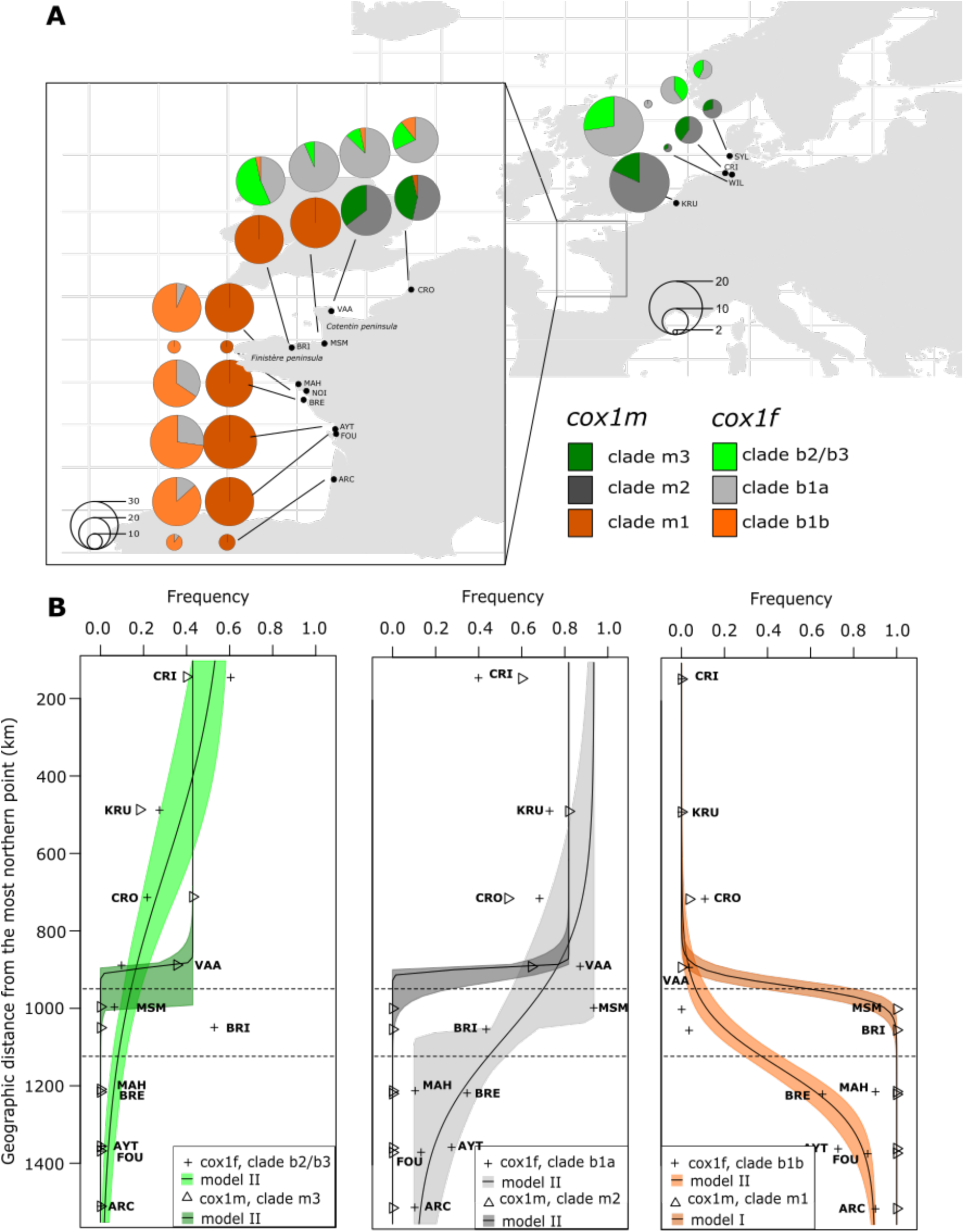
Geographical distribution of the *cox1f* and *cox1m* haplogroups: (A) haplogroup frequencies are represented for each marker in all the sites with a pie chart. The size of the pie chart indicates the number of individuals analyzed, (B) results of the cline analyses is presented with comparison between male- and female-type haplogroup. The transparent colored area around the fitted curve represents the 95% confidence interval. The haplogroup m4 is not presented here as it is only found in one individual from GRI.

Saint-Brieuc (BRI) for instance was highly differentiated from all other northern sites except from MSM (Mont-Saint-Michel). Based on these results, genetically homogenous sampling sites were merged in 3 geographical groups (Table 1) and used in subsequent analysis (North, Center and South). Sampling sites within the hybrid/Finistère group showed genetic differentiation only for *cox1f*.

Cline analyses performed on haplogroup frequencies consistently revealed clinal variations with null model rejected for each haplogroup (Figure 2B and Table S5). For the major northern clades (b1a/m2) and the southern clades (b1b/m1), a geographical discordance in the center of hybrid zone was observed, ranging from 178 (clades b1a vs. m2) to 209 (clades b1b vs. m1) km. In both cases (major northern clades and southern clades), the genetic break was located on the Cotentin Peninsula for *cox1m*, and the western tip of the Finistère Peninsula for *cox1f*. The width estimated from the maximum likelihood cline of the best-fit models were 4 to 33 times narrower at *cox1m* (16 to 80 km), compared to *cox1f* (317 and 540 km for clades b1b and b1a, respectively). Concerning the minor northern clades (b2/b3 and m3), the null model was rejected but the clinal model might not be the best fit to explain its distribution. Indeed, b2/b3 was absent in the south but minor in the northern and central sites (from 6 to 27%), except for CRI and BRI where it was dominant. BRI actually accounts for a large part of b2/b3’s diversity, with 5 of the 10 haplotypes in this group, four of which are private to this sampling site. This geographic pattern contrasts with m3, which followed a clinal model: a frequency ranging between 18 and 42% in the north and absent in the south.

Haplotypes from the extreme haplogroups (S and N) were rarely shared within one individual (Figure 1B), and while 17 individuals exhibited the haplogroup b2/b3 at *cox1f* and the haplogroup m1 at *cox1m*, only 3 individuals shared the *cox1f* haplogroup b1b and the *cox1m* haplogroup m3. Also, only 1 individual presented both the *cox1f* haplogroup b1b and the *cox1m* haplogroup m2, suggesting non-random mating or lethal haplotype combinations. Noteworthily, most of these individuals were found in the hybrid zone sampling sites (18 out of 22). These results were confirmed by the significant linkage disequilibrium test carried out among haplogroups for phased data in the hybrid zone and the south (T_2 test df=6, p<0.001).

The level of genetic diversity also differed between the two markers (Table 1 and Figure 3). While the average nucleotide diversity (p) and its range across sampling sites were similar (a mean value of 0.0047 and a range of 0 to 0.0091 for *cox1f*; a mean of 0.0019 and a range 0.0004 to 0.007 for *cox1m*), the median values of the nucleotide diversity (0.38 and 0.12 for *cox1f* and *cox1m* respectively) showed that the nucleotide diversity is significantly greater for *cox1f* than for *cox1m* (Wilcoxon rank test V= 65 p= 0.002). We observed the lowest nucleotide diversities in southern sampling sites for *cox1m* compared to *cox1f*. The haplotype diversity was not significantly different between the two markers (Wilcoxon rank test, V = 35, p = 0.89). Finally, Tajima’s D also differed among the two markers. When considering the three geographical groups, significant negative values were found only for *cox1m* indicating the possibility of population expansion after a bottleneck event or selective sweep. For *cox1f*, on the other hand, no sign of deviation from neutral expectations was detected at the population level. The proportion of singletons at the scale of sample sites (Figure 3) showed an overall higher number of rare frequency mutations in *cox1m* compared to *cox1f*, which combined with negative D values, could indicate a recent expansion in population size. Mismatch distribution analyses and drift-mutation equilibrium tests suggest different demographic histories at *cox1f* and *cox1m*, with bimodal pairwise nucleotide distributions at the former, and unimodal right-skewed distributions at the latter (Figure S4 and Table S5). Based on the non-significant raggedness indices (r), for both markers and groups, the hypothesis of sudden population expansion cannot be rejected; this result is only substantiated at *cox1m* (in both groups), by the R_2_ statistic. This index estimated as the difference between the number of singleton mutations and the average number of nucleotide difference among haplotypes has been shown as the most powerful to detect population expansion in case of small sample size and few segregating sites (Ramos-Onsins & Rozas, 2002). While molecular signatures of the mutation/drift (dis)equilibrium are similar between southern and northern groups regardless of the marker, the mismatch distributions show more divergence among haplotypes in the North (pairwise nucleotide differences >6), compared to the South (pairwise nucleotide differences <6).

**Figure 3.**
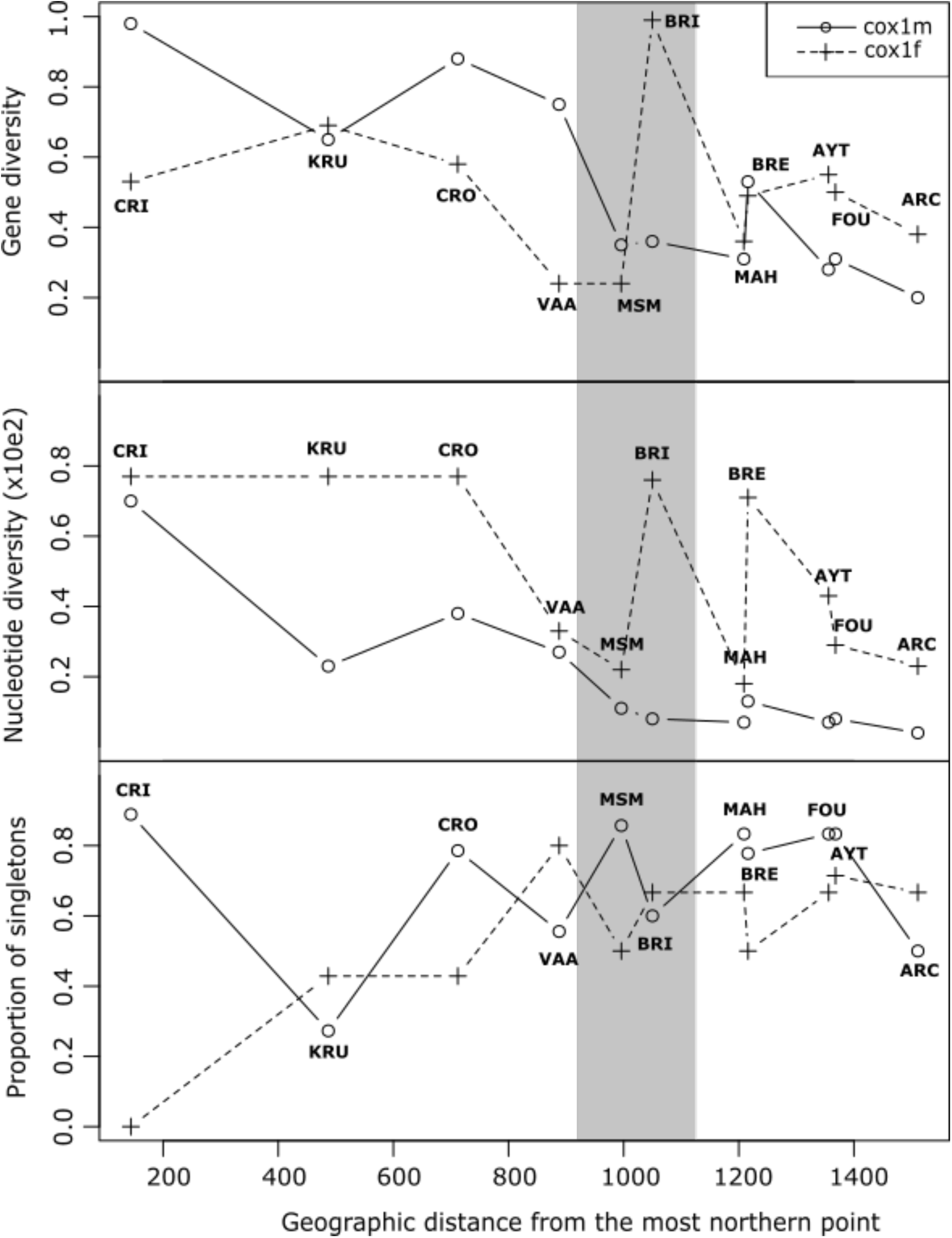
Genetic diversity indexes for both markers (*cox1m* and *cox1f*) along the coast. The gray area indicates the geographic range between the Cotentin and the Finistère Peninsulas.

Consistently with the haplotype networks, the results of the AMOVAs suggested different geographic structuration patterns for *cox1m* and *cox1f* (Table 2). The best hierarchical model based on the F_SC_/F_CT_ ratio (least variation within and highest between groups) was different for the two markers: for *cox1f* the best model was two groups (B) separated by the Finistère peninsula (KRU, CRI, CRO, VAA, MSM, BRI *vs* MAH, BRE, AYT, FOU, ARC) whereas for *cox1m*, the best model was also two groups of subpopulations but (A) with the Cotentin peninsula as a major geographical break (KRU, CRI, CRO, VAA) *vs* (MSM, BRI, MAH, BRE, AYT, FOU, ARC). The result of the AMOVAs (Table 2) showed similar level of among subpopulation genetic variation relative to the total variation for both markers (*cox1m*: 42% and Φ_ST_= 0.58, p<0.001; *cox1f*: 44% and Φ_ST_= 0.55, p<0.001). The level of differentiation among groups was higher for *cox1m* compared to *cox1f* (Φ_CT_= 0.57, p<0.001 and Φ_CT_= 0.49, p<0.001 for *cox1m* and *cox1f*, respectively). Finally, within group genetic structure was detected only for *cox1f* (Φ_SC_= 0.01, p=0.062 and Φ_SC_= 0.13, p<0.001 for *cox1m* and *cox1f*, respectively). Genetic differentiation analyses based only on haplotype frequencies (conventional F-statistics) were also carried out and gave similar results (not shown).

**Table 2.** Analysis of molecular variance (AMOVA) to partition the genetic variation among hierarchical geographical scales. Results are presented for both markers, with different options of hierarchical grouping of subpopulations.

| Number of groups | Groups |  | *Cox1f* |  |  | *Cox1m* |  |  |
| --- | --- | --- | --- | --- | --- | --- | --- | --- |
|  |  |  | Among groups relative to total variance | Among subpopulations within groups | Among subpopulations relative to total variance | Among groups relative to total variance | Among subpopulations within groups | Among subpopulations relative to total variance |
| 3 | (CRI,KRU) (CRO,VAA,MSM,BRI)<br>(MAH,BRE,AYT,FOU,ARC) | %var | 44.71 | 7 | 48.29 | 22.2 | 22.54 | 55.26 |
| | | Fixation indices | $\Phi_{CT}=0.447^{***}$ | $\Phi_{SC}=0.517^{***}$ | $\Phi_{ST}=0.126^{**}$ | $\Phi_{CT}=0.221^{ns}$ | $\Phi_{SC}=0.290^{***}$ | $\Phi_{ST}=0.447^{***}$ |
| 2(A) | (CRI,KRU,CRO,VAA)<br>(MSM,BRI,MAH,BRE,AYT,FOU,ARC) | %var | 16.98 | 31.02 | 52 | 57.36 | 0.56 | 42.08 |
| | | Fixation indices | $\Phi_{CT}=0.170^{ns}$ | $\Phi_{SC}=0.374^{***}$ | $\Phi_{ST}=0.480^{***}$ | $\Phi_{CT}=0.574^{***}$ | $\Phi_{SC}=0.013^{ns}$ | $\Phi_{ST}=0.579^{***}$ |
| 2(B) | (CRI,KRU,CRO,VAA,MSM,BRI)<br>(MAH,BRE,AYT,FOU,ARC) | %var | 49.14 | 6.4 | 44.46 | 21.88 | 4.18 | 53.94 |
| | | Fixation indices | $\Phi_{CT}=0.491^{***}$ | $\Phi_{SC}=0.126^{***}$ | $\Phi_{ST}=0.555^{**}$ | $\Phi_{CT}=0.212^*$ | $\Phi_{SC}=0.309^{***}$ | $\Phi_{ST}=0.460^{***}$ |

Note: \*P>0.05, \*\*P>0.01, \*\*\*P < 0.001, ns: non significant

Finally, our results suggest that either Ne and/or μ differed geographically. The theta estimates were twice larger at *cox1m* compared to *cox1f* in the North (Theta_f/Theta_m =0.48) and similar in the South (Theta_f/Theta_m=0.98).

## Discussion

### Phylogeography of *Macoma* lineages and F/M geographic discordance

Recovering the M- and F-type *cox1* of *M. b. balthica* from the transcriptomic study of Yurchenko et al., (2018) allowed the application of the net divergence method used in Nikula et al., (2007) on both *cox1f* and *cox1m* datasets. As detailed above, this clock hypothesizes that *rubra* and *balthica* diverged between 3.5-2 Mya (bounded by the opening of the Bering Strait and the discovery of first Atlantic fossils, respectively) and was applied by Nikula et al., (2007) on *cox3f*. At the scale of the genus, *cox1f* and *cox1m* data confirmed that the divergence between *M. balthica* and *M. petalum* occurred at the end of the Miocene, early Pliocene at the latest (*cox1f*: 4.9-8.6 My; *cox1m*: 5.4-9.5 My), as inferred with *cox3f* (5.2-9.3 My; Nikula et al., 2007). Within *M. balthica*, estimates at *cox1f, cox1m* and *cox3f* congruently suggest a split between clades C and D during the Calabrian subdivision of the Pleistocene (0.83-1.76, 0.98-1.72, 0.8-1.4 My, respectively), again confirming previous results (Nikula et al., 2007).

In the North (Crildumersiel, Germany) and Baltic Seas (Mecklenburg bight, Germany; Lomma, Sweden, in the Öresund Strait; Umea, Sweden, in the Gulf of Bothnia), a distinct *M. b. rubra* clade here called m4 was sampled at *cox1m* (Table S2, Figure S2). This clade diverged from m3 about 220 to 380 kya and is probably the M-type counterpart of b2/b3, which diverged from b1a about 251 to 440 kya. Males with m4 haplotypes possessed either b2/b3 or D at *cox1f*. The b2/b3 clade is common in North Sea, the Öresund and Western Baltic (Nikula et al., 2007), making its association with m4 expected. However, six m4 specimens were sampled from Umea, five of which bearing a *M. b. balthica* signature at *cox1f* (clade D), showing that *rubra* m4 introgressed far north into the Baltic Sea, as does b2/b3 (Nikula et al., 2007).

The split between the southern and northern *M. b. rubra* clades, currently separated by the Gulf of Saint-Malo where they hybridize, occurred much more recently. At *cox1f*, divergence was estimated between 106 and 185 kya, in opposition with the much later split estimated at *cox1m* (30-50 kya). These different split times may be due to the different forces acting upon the M- and F-type mtDNA, namely the selection / mutation equilibrium (known to differ between M- and F-type mtDNA, see below), drift (which may have been substantial during glacial cycles), assortative mating (a pre-zygotic barrier, see Bierne et al., 2002 for an example in *Mytilus*), sex-biased migration (see Arnaud-Haond et al., 2003 for an example in oysters), and different hybrid fitness between sexes. Nevertheless, both split times are consistent with a scenario of pre-LGM vicariance (the LGM is currently estimated to have occurred between 26.5 and 19 kya; Clark et al., 2009), in opposition to a post-glacial scenario of primary intergradation.

Zooming in on the Gulf of Saint-Malo hybrid zone revealed discordant phylogeographic patterns for the female- and male-type mitochondrial *cox1* gene. Both male and female mtDNAs displayed significant segregation of haplotypes in space. As previously described by Becquet et al., (2012), a geographical break at the Finistère peninsula was observed at *cox1f*. Sampling done for the present study narrows the gap between northern and southern populations by about 50 km thanks to the addition of the site of Saint-Brieuc Bay, however we failed at sampling any large population between the latter and Pont-Mahé Bay, south of Finistère (Figure 2). While anecdotal data support the presence of *M. b. rubra* in the Bay of Brest (Hily, 2013 and our observations), the Finistère intertidal seem largely inhospitable to this species due to the scarcity of mud flats (Becquet et al., 2012, Mao et al., 2020 and this study). It therefore appears as a credible physical barrier to genetic connectivity, and it has been indeed identified in the past as a genetic breaking zone for other marine benthic-pelagic species confined to estuaries and bays (muddy-fine sediment species, Jolly et al., 2005, 2006).

On the other hand, the Cotentin peninsula located about 250 km farther East of the Finistère peninsula separated southern and northern populations *cox1m* haplotypes at a much narrower hybrid zone (an observation that might be interpreted by a different selection / migration balance compared to *cox1f*, or the heterogeneous spacing of sampling points along the coast). Sandy mud flats are numerous along western Cotentin but do not seem hospitable to *M. balthica.* Our field surveys failed at identifying populations between Mont-Saint-Michel Bay (MSM) and Saint-Vaast-la-Hougue (VAA), although anecdotal presence of the species was recorded farther along the Cotentin coast and was reported to be present but rare on the Gulf of Saint-Malo (Le Mao et al., 2020). Along with the scarcity of seemingly appropriate habitat and patchiness of small populations in the intertidal zone of Cotentin, we observed significant differences in gonadal development and spawning phenology between populations on either side of the peninsula. While individuals with full gonads and well-developed gametes were sampled between the 4th and the 23rd of April 2018 from Fouras to Mont-Saint-Michel Bay (some individuals having even spawned in the field at MSM and Saint-Brieuc), gonads from Saint-Vaast-la-Hougue (VAA) and Somme Bay individuals were still in early stages of development by April 19, 2018 (most gametes being still undifferentiated). These observations were on par with previous reports suggesting spawning asynchrony across the Cotentin Peninsula, with spawning occurring in early April in Aytré (Saunier, 2015) and August in Somme Bay (Ruellet, 2013). In the great scallop, *Pecten maximus*, different spawning phenologies were described across the Cotentin peninsula (Lubet et al., 1991). As for the Finistère, the Cotentin peninsula was previously identified as a barrier to gene flow (Jolly et al., 2005; Quéméré et al., 2016; Handal et al., 2020). It is also the southern biogeographical boundary of the northern cold temperate Boreal region (Dinter, 2001) with distinct hydrologic and oceanographic features from the western part of the English Channel (Dauvin, 2012). There are, therefore, multiple exogenous (habitat availability; oceanographic currents, *e.g.* Dupont et al., 2007; Fievet et al., 2007; Hily, 2013) and endogenous (asynchrony in reproductive phenology and genetic incompatibilities, discussed below) barriers that could contribute to the genetic differentiation of populations across these two peninsulas.

The fact that the geographic discordance of genetic clines is sex-specific is quite original for a marine invertebrate. In the DUI literature, studies have reported that genetic differentiation across populations is stronger for M-type mtDNA than for F-type mtDNA (Liu et al., 1996; Riginos et al., 2004; Skibinski et al., 1999), and some authors reported discordant genetic structure between markers. The freshwater mussels *Lampsilis siliquoidea* displayed population genetic structure at *cox1f* but none at *cox1m* (Krebs et al., 2013). Conversely, Riginos et al., (2004) found weak connectivity at F-type mtDNA but none at M-type mtDNA in *Mytilus edulis* across the Atlantic Ocean. At the entrance of the Baltic, rampant introgression of *M. edulis* F-type mtDNA was observed, compared to a sharp cline at M-type mtDNA concordant with M7 lysin (a nuclear gene involved in fertilization; Stuckas et al., 2009). This makes the population structure described here for *M. b. rubra* noteworthy in the sense that the phylogeography of F- and M-type mtDNA markers do not simply differ in amplitude or resolution (due to significant differences in mutation rates, μM being about twice μF) but differ in the geographical position of the haplotypic cline separating southern and northern *M. b. rubra*.

While a genetic break located around the Finistère peninsula corresponds to previous results by Becquet et al., (2012) for *cox1f* and eight nuclear microsatellites, the Cotentin was not detected as a barrier in that study. However, spatial structure was detected at the nuclear *atp5c1* gene (encoding the gamma subunit of the FO/F1 ATP synthase protein complex) between Mont-Saint-Michel and Somme Bay (Saunier, 2015). Non-synonymous point mutations separating southern and northern populations are located in an inter-peptide interaction domain of the gamma subunit, suggesting mito-nuclear incompatibilities (Pante et al., 2019). The concordance of *atp5c1* and *cox1m* calls for further investigation of incompatibilities involving the male mitogenome specifically. Interestingly, significant differentiation with *cox1f* was detected between Pont-Mahé Bay (MAH) and the Saint Brévin (BRE) and Aytré (AYT) populations (Table S4). These two groups, separated by the Vilaine River, were identified as significantly differentiated based on microsatellites, but not *cox1f* in the study of Becquet et al., (2012), which was based on individuals from both sexes, while our study focused on males exclusively. We may therefore benefit from higher statistical power to detect differences in differentiation across the Vilaine River. This implies that genetic structure among males at *cox1f* is higher than in females. Although we did not include females in our study, we did compare pairwise *Φ*_ST_ values between sampling sites that were common between our study and Becquet’s (2012). *F*_ST_ and *Φ*_ST_ were both significantly higher in the dataset composed of males exclusively, compared to the dataset composed of males and females (Figure 4). This pattern can be attributed to stochastic genetic variation (effects of mutations and drift) and/or differences of gene flow between sexes. The geographical discordance observed at the Finistère and Cotentin peninsulas for *cox1f* and *cox1m* are also reminiscent of sex-level differences. Indeed, in studies using female-transmitted mtDNA and male-transmitted Y chromosome to investigate population structure, sex-biased asymmetries are often cited as a determinant of discordant geographic patterns (*e.g.* Boissinot & Boursot, 1997; Trejo-Salazar et al., 2021). Sex-level differences in gene flow are scarcely reported but exist in bivalves, as in the protandrous pearl oyster *Pinctada mazatlanica*, which effective sex-ratio is strongly biased towards males (Arnaud-Haond et al., 2003).

**Figure 4.**
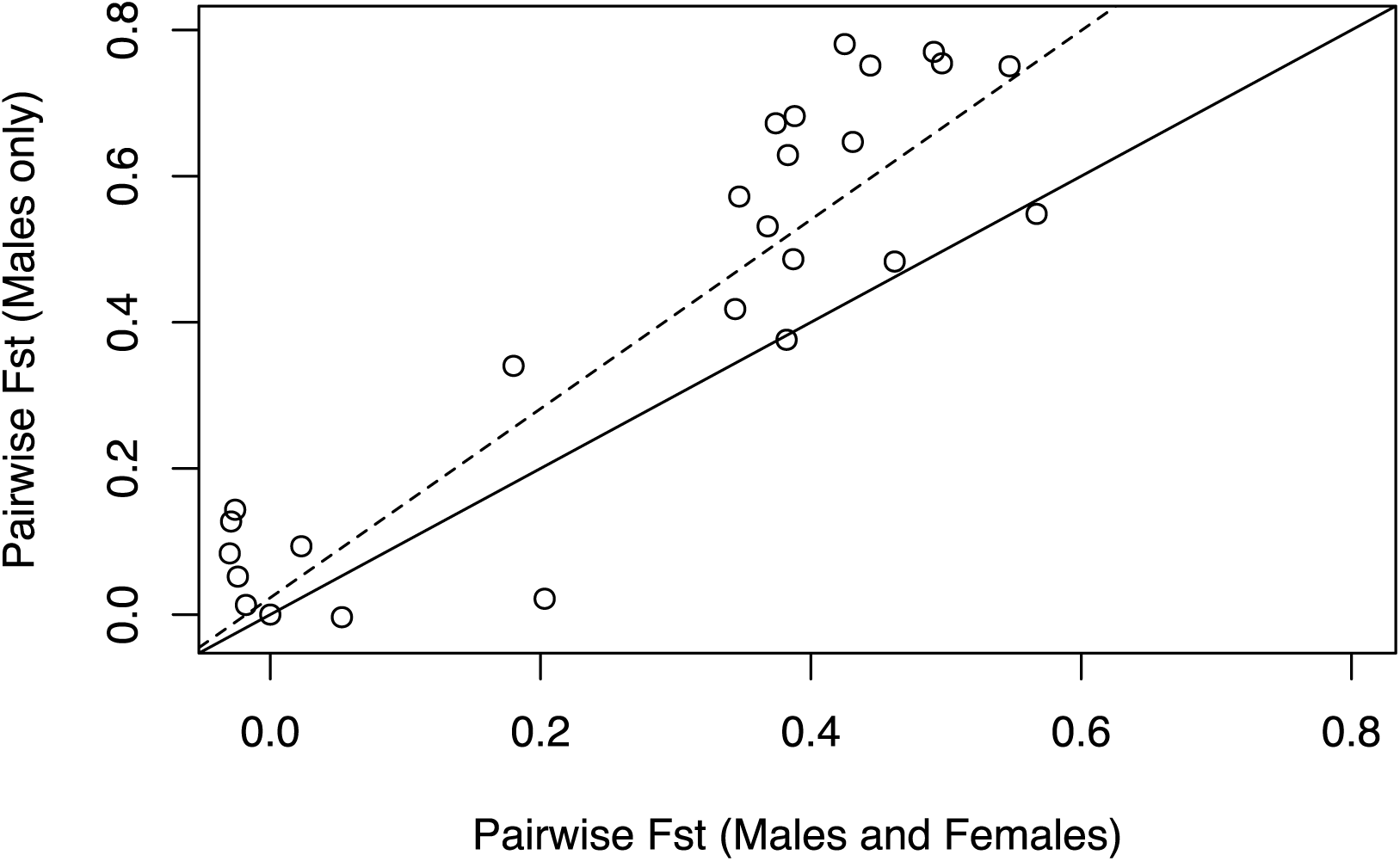
Comparison of pairwise F_ST_ estimated at the *cox1f* gene for study sites that were both sampled by Becquet et al (2012) (male and female individuals) and for the present study (male individuals only). The plain line represents the x=y axis and the dash line, the linear regression (y = 1.27 x + 0.03, df = 62, R^2^_adj_=0.74, p<0.001). A Mantel test revealed a significant correlation between the two datasets (r=0.786, p=0.0084).

Other factors could explain the M- vs. F-mtDNA discordance, such as mtDNA introgression and heterogeneity in its rate, hybrid zone spatial instability and drift (Barton & Hewitt, 1985; Bierne et al., 2003; Toews & Brelsford, 2012). Patterns of asymmetric introgression of alleles are commonly observed in hybrid zones and were observed in the Bay of Biscay between *Mytilus edulis* and *M. galloprovincialis* (Rawson & Hilbish, 1995). In *M. b. rubra*, asymmetrical gene flow from North to South was detected at *cox1f* (Becquet et al., 2012, this study), and *atp5c1* (Saunier, 2015; Pante et al., 2019). At that same locus, Gagnaire et al., (2012), looking at divergence and selection among populations of the eels *Anguilla rostrata* and *A. anguilla* (two hybridizing sister species) also found evidence of unidirectional introgression from the former to the later. At *cox1m*, little introgression was detectable in *M. b. rubra*, but nevertheless sensibly higher from north to south. Hybrid zone movement has been recognized for long, as a manifestation of variation in population density, dispersal rate, individual fitness, effects of allele frequencies on population structure, and spatially- or frequency-dependent selection (Barton & Hewitt, 1985). It has been proposed that the northeastern Atlantic *Mytilus* hybrid zones move parallel to warming sea surface temperatures (Hilbish et al., 2012). While a similar phenomenon may occur in Baltic tellins, suggesting that genotype-environment correlations could affect cline geography (Ducos et al unpublished), neutral forces may be sufficient to result in discordant M-type and F-type mtDNA cline centers and width. Indeed, as detailed above, both the Finistère and the Cotentin peninsulas are characterized by low population densities and oceanic currents disrupting along-coast dispersal. While mitochondrial recombination is expected to be rare (or at least, rarely transmitted to offspring; Passamonti et al., 2003), looking at whole mitogenomes as well as nuclear genes of mitochondrial function vs. other (putatively neutral) nuclear markers may help shed light on the degree to which the patterns detected at *cox1* and *atp5c1* depart from neutral expectations.

Whether the asymmetric discordant clines in *M. balthica* observed here are due to cline movement, differential gene flow, population structure, or selection (Barton & Hewitt, 1985) remains a fascinating avenue for future research. In particular, introgression across the hybrid zone in the English Channel calls for a test of asymmetric fitness of inter-lineage crosses (e.g., Turelli & Moyle, 2007). An involvement of mito-nuclear incompatibilities could explain the observed asymmetric allele frequencies if northern hybrids with southern mitochondria are less fit than southern hybrids with northern mitochondria.

### Genetic diversity differences between *cox1m* and *cox1f*

F- and M-mtDNA diversity was compared among homologous regions within heteroplasmic males, and therefore variability can be strictly attributed to heredity rather than inter-individual or inter-genic differences. M-type mtDNA was almost up to twice as polymorphic as its female counterpart, with more haplotypes, more singletons (*i.e.* haplotypes represented by a single male) and more segregating sites. Higher polymorphism in the M-type mtDNA, both within and between species (Skibinski et al., 1999), is commonly reported in DUI species, as in *Polititapes rhombroides* (Chacón et al., 2020), *Pyganodon grandis* (Krebs, 2004) and *Mytilus sp.* (Ort & Pogson, 2007; Śmietanka et al., 2009, 2013; Śmietanka & Burzyński, 2017). Higher frequency of rare M-type haplotypes (80% of singletons compared to 57% among F-type haplotypes in *M. b. rubra*) is another largely shared feature among DUI species (Ort & Pogson, 2007; Śmietanka et al., 2009, 2013; Śmietanka & Burzyński, 2017; Chacón et al., 2020). Major determinants of genetic diversity in animals are linked selection, mutation, and effective population size (Ellegren & Galtier, 2016). In DUI bivalves, polymorphism might result from different selection pressures on male- and female-type mitogenomes (*e.g.* Milani et al., 2012): female-type mitochondria ensure proper mitochondrial function at the scale of the individual whereas male-type mitochondria, which solely co-occurs in heteroplasmic individuals with the F-type mitochondria (Skibinski et al., 1994; Zouros et al., 1994) have to effectively function exclusively in the male germ line and gametes (different ‘arenas of selection’ *sensu*, Stewart et al., 1996). According to the male-driven evolution hypothesis, higher germ cell divisions in males (Shimmin et al., 1993) could result in an enhanced replication rate of the M-type mitochondria during spermatogenesis, providing more opportunities for mutations to accumulate (Rawson & Hilbish, 1995; Stewart et al., 1995). In addition, oxidative damage might be important in sperm (Zouros, 2013), and/or selection pressures on chaperonins and DNA repair genes might be relaxed. We can also expect sex-linked demographic differences to influence polymorphism, if the effective population size differs between males and females; we did observe larger theta at *cox1m* in northern populations, suggesting that the effective population size and/or mutation rate is larger in males. Most hypotheses on the role of selection, mutation and effective population size in shaping genetic diversity in DUI bivalves remain to be empirically tested.

However, counter examples do exist. For instance, Beauchamp et al., (2020) found more F-type haplotypes than M-type haplotypes in *Pyganodon grandis* (11 and 7, respectively) and *P. lacustris* (20 and 6, respectively). Śmietanka et al., (2013) (87 F- and 76 M-type haplotypes in *Mytilus trossulus*) and Riginos et al., (2004) (132 F- and 56 M-type haplotypes in *Mytilus edulis*) reported the same pattern, although the latter results might be due to the difficulty of amplifying the male-type mtDNA. Although often presented as such in the literature, higher polymorphism in males is therefore not a universal feature of the population genetics of DUI species.

It is worth noting that, among the studies reporting M/F differences in genetic polymorphism, all of those that explicitly tested for non-equilibrium population state found support for population expansion (Chacón et al., 2020; Ort & Pogson, 2007; Riginos et al., 2004; Śmietanka et al., 2009, 2013; Śmietanka & Burzyński, 2017, and our study at *cox1m*), except for one finding support for purifying selection (Skibinski et al., 1994). Non-equilibrium states following a glacial period can indeed affect genetic diversity, notably through drift, and confound studies focusing on the comparative molecular evolution of F and M mtDNA. It would therefore be interesting to look for biological models at equilibrium.

## Data Accessibility

*Cox1f* and *cox1-long* sequences were submitted to Genbank (accession N° OM855617 - OM855929 and OM856027 - OM856339 respectively). Rcode is available at https://github.com/SabLeCam/Cox1_DUI

## Acknowledgements

The authors thank Mélanie Rocroy, Patrick Triplet, Thierry Ruellet, Rolet Céline (Association GEMEL, Picardie), Anthony Sturbois (RNN Baie de Saint-Brieuc, VivArmor Nature) and Xavier de Montaudouin (Station Marine d’Arcachon, UMR 5805 EPOC, Université de Bordeaux/CNRS) for their help with sample collection. We thank the participants of the Biocombe project (“The impact of BIOdiversity changes in COastal Marine Benthic Ecosystems;” contract n. EVK3-CT-2002-00072; Research n. OND1290723). Lab work was performed at the Molecular Core Facility at the University of La Rochelle. We thank the bioinformatics platform Toulouse Midi-Pyrénées (Bioinfo GenoToul). Preprint version 4 of this article has been peer-reviewed and recommended by Peer Community In Evolutionary Biology (https://doi.org/10.1101/2022.02.28.479517; Castilho, 2025.

## Data, scripts, code, and supplementary information availability

Data, scripts, and supplementary information are available online: https://doi.org/10.5281/zenodo.14849342

