## supplementary material for "Discordant population structure inferred from male- and female-type mtDNAs from *Macoma balthica*, a bivalve species characterized by doubly uniparental inheritance of mitochondria" (from another version of this preprint: version 4, 2025-01-17)

Table S1 Sampling sites, ID, geographic coordinates (WGS84) and the country of the locality.

| **Species** | **Sampling site** | **ID** | **Latitude** | **Longitude** | **Country** |
| --- | --- | --- | --- | --- | --- |
| *Macoma balthica* | Vancouver | VANC | 49.07 | -123.18 | Canada |
|  | Departure Bay | D.BAY | 49.06 | -123.97 | Canada |
|  | Baie Comeau | H5M,G5M | 49.16 | -68.19 | Canada |
|  | Umea | UME | 63.83 | 20.24 | Sweden |
|  | Lomma | LOM | 55.68 | 13.07 | Sweden |
|  | Mecklenburg bight | MEC | 53.90 | 12.5 | Germany |
|  | Sylt | SYL | 54.90 | 8.38 | Germany |
|  | Wilhelmshaven | WIL | 53.54 | 8.57 | Germany |
|  | Crildumersiel | GRI | 53.65 | 8.02 | Germany |
|  | Kruiningen | KRU | 51.45 | 4.04 | Netherlands |
|  | Le Crotoy | CRO | 50.21 | 1.62 | France |
|  | Saint-Vaast-la Hougue | VAA | 49.58 | -1.26 | France |
|  | Mont-Saint-Michel | MSM | 48.63 | -1.51 | France |
|  | Saint-Brieuc | BRI | 48.51 | -2.69 | France |
|  | Pont-Mahe | MAH | 47.44 | -2.45 | France |
|  | Saint-Brévin | BRE | 47.24 | -2.16 | France |
|  | Noirmoutier | NOI | 46.98 | -2.26 | France |
|  | Aytre | AYT | 46.12 | -1.12 | France |
|  | Fouras | FOU | 45.98 | -1.09 | France |
|  | Arcachon | ARC | 44.65 | -1.17 | France |
| *Macoma petalum* | Addison | ADD | 44.62 | -67.74 | United States |
|  | York River | YR | 37.41 | -76.71 | United States |

| Table S2 Net distance (da) between monophyletic *Macoma* lineages and estimates of cladogenesis age. Upper estimates are based on the date of the earlier *M.balthica* (2 My) and lower estimates , on the opening of the Bering strait (3.5 My). | ***cox1f*** | | | |  | ***cox1m*** | | | |
| --- | --- | --- | --- | --- | --- | --- | --- | --- | --- |
| **Contrast** | **Clades** | **da (s/s)** | **upper (My)** | **lower (My)** |  | **Clades** | **da (s/s)** | **upper (My)** | **lower (My)** |
| *M.petalum - M. balthica* | A vs bi/C/D | 0,217 | 4,898 | 8,571 |  | A vs mi/C/D | 0,447 | 5,40 | 9,46 |
| *M. b. rubra- M. b. balthica lineages* | bi vs C/D | 0,089 | 2,000 | 3,500 |  | mi vs C/D | 0,165 | 2,00 | 3,50 |
| *M. b. balthica groups* | C vs D | 0,037 | 0,833 | 1,459 |  | C vs D | 0,0814 | 0,98 | 1,72 |
| *M. b. rubra groups* | b1a vs b2/b3 | 0,011 | 0,251 | 0,440 |  | m1 vs m4 | 0,022 | 0,26 | 0,46 |
| *M. b. rubra groups* | b1b vs b2/b3 | 0,017 | 0,386 | 0,676 |  | m2 vs m4 | 0,020 | 0,24 | 0,42 |
| *M. b. rubra groups* | b1a vs b1b | 0,005 | 0,106 | 0,185 |  | m3 vs m4 | 0,018 | 0,22 | 0,38 |
| *M. b. rubra groups* | -- | -- | -- | -- |  | m1 vs m2/m3 | 0,002 | 0,03 | 0,05 |

Table S3 Pairwise Φ_ST_ values among population based on *cox1m* (significant values in bold based on 10 000 permutations).

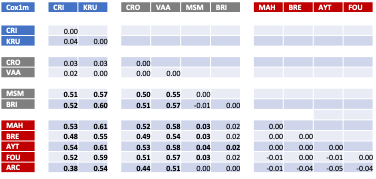

Table S4 Pairwise Φ_ST_ values among population based on *cox1f* (significant values in bold based on 10 000 permutations).

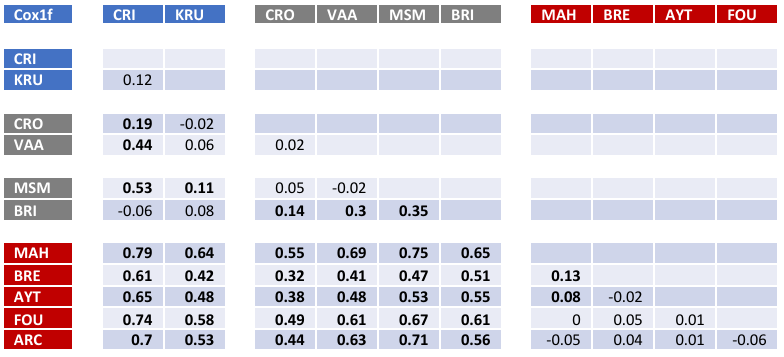

Table S5: Results of the cline analyses on haplogroup frequency. The best-fit model is indicated together with the AIC score difference with the null model and the model parameters (center and width of the cline).

| **Marker** | **Halogroup** | **Best Model** | **ΔΑΙC with Null model** | **Center (km)** | **Width (km)** |
| --- | --- | --- | --- | --- | --- |
| *Cox1f* | Clade b2/b3 | II | 30.14 | 641 | 1052 |
|  | Clade b1a | II | 45.68 | 1071 | 540 |
|  | Clade b1b | II | 169.67 | 1145 | 317 |
| *Cox1m* | Clade m3 | II | 74.80 | 895 | 18 |
|  | Clade m2 | II | 160.62 | 893 | 16 |
|  | Clade m1 | I | 312.70 | 936 | 80 |

Model types : Null (no cline), I (fixed scaling based on minimum and maximum observed frequencies and no exponential tails) and II (free scaling and no exponential tails).

Table S6: Harpending's raggedness (r), Ramos-Onsins & Rozas' (R_2_), Fu's Fs and Tajima’s D estimates for the mismatch distributions are presented together with upper (UL)and lower (LL) confidence interval (95%), standard deviation (S.D.) and p value ( based on 10 000 permutation tests)

|  |  | **r** |  | **R_2_** |  | **Fs** |  | **D** |  |
| --- | --- | --- | --- | --- | --- | --- | --- | --- | --- |
| Cox1f | North | 0.1878 | LL 0.01971  UL: 0.32734  p=0.89190 | 0.1423 | LL: 0.05663  UL: 0.18384  p=0.82078 | 1.78412 | p= 0.87580 | 0.75250 | S.D=1.10663  p=0.71910 |
|  | South | 0.1960 | LL: 0.02656  UL 0.56222  p=0.74705 | 0.0538 | LL: 0.03592  UL: 0.17487  p=0.14065 | **-5.86889** | p=0.01950 | -1.14611 | S.D.=1.11794  p=0.09620 |
| Cox1m | North | 0.0730 | LL 0.02930  UL 0.62635  p=0.23072 | **0.0553** | LL: 0.05720  UL: 0.20054  p=0.01855 | **-10.94649** | p=0.00000 | **-2.25856** | S.D.=1.12402  p=0.00060 |
|  | South | 0.1835 | LL: 0.07513  UL: 0.92349  p=0.25714 | **0.0232** | LL: 0.01941  UL 0.24073  p=0.03563 | -**22.60307** | p=0.00000 | **-2.40393** | S.D.=1.12976  p=0.00000 |

Figure S1: Maximum-likelihood phylogenetic tree of *Macoma balthica* *cox1f* haplotypes and the outgroup *Macoma petalum*. Haplotypes signaled with a circle are from Nikula et al. 2007 (Accession numbers: EF0044-125,127:130,132,136), stars from Becquet et al. 2012 (Accession numbers: HM756170-89), squares from Layton et al. 2016 (Accession numbers: KP775-84,87,92; KP776-15,21,32,38; KP9777-09,12,50,54,70,84; KP9778-10,21,43,82; KP9779-36,58,61) and triangle from Yurchenko et al. 2018 (Accession project number: PRJNA384460) . Bootstrap support (out of 1000 >80%) is indicated at each branch. Locations of the samples are presented in figure S3.

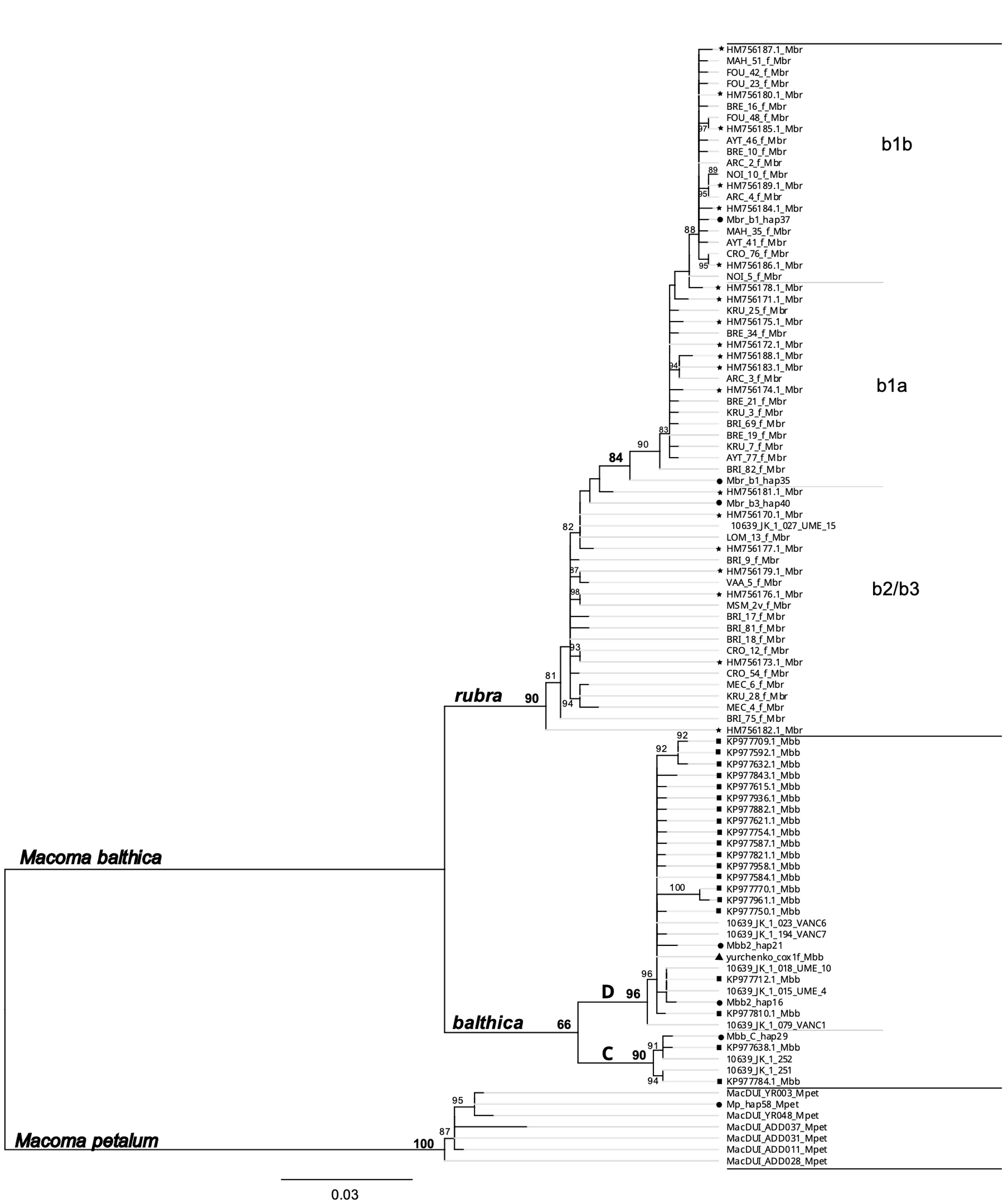

Figure S2: Maximum-likelihood phylogenetic tree of *Macoma balthica* *cox1m* haplotypes and the outgroup *Macoma petalum.* Tips with asterisks are m2 haplotypes grouped with m1. Haplotypes signaled with a star are from Yurchenko et al 2018 (Accession project number: PRJNA384460). Bootstrap support (out of 1000 > 80%) is indicated at each branch. Locations of the samples are presented in figure S3.

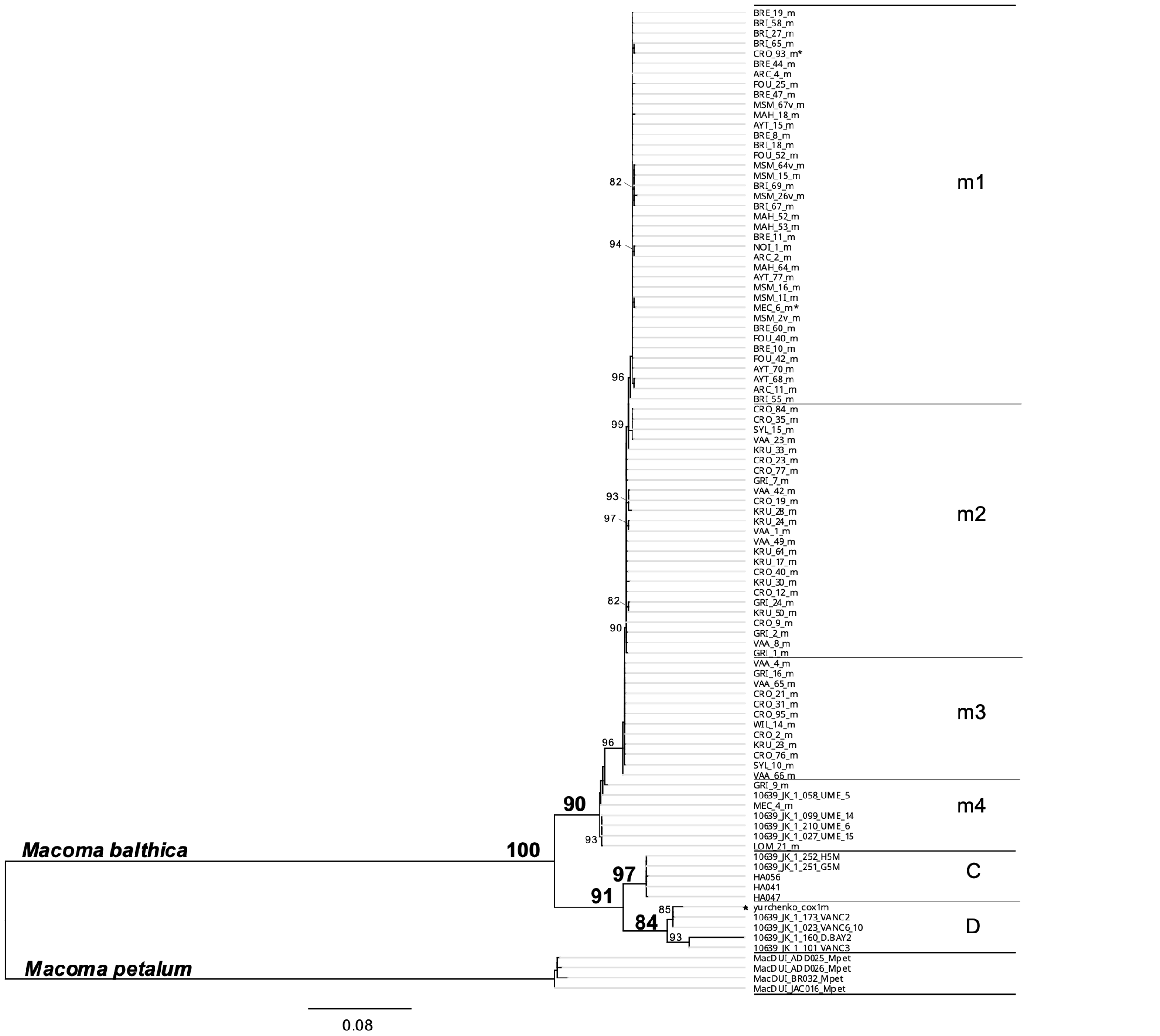

Figure S3: Map of samples considered in the phylogenetic trees in figures S1 and S2

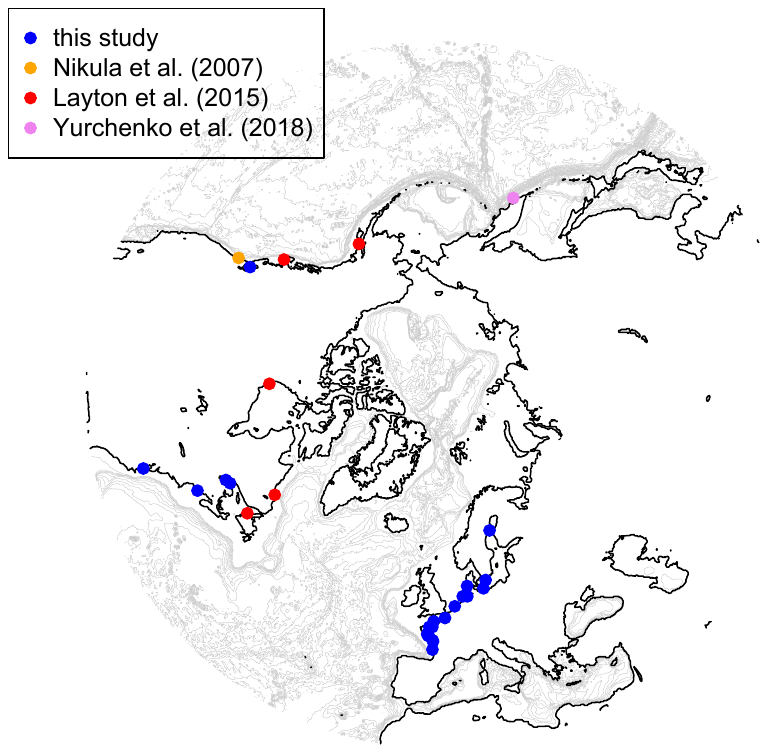

Figure S4: Mismatch analyses of North and South populations at both markers. Histograms represent the observed data of pairwise differences; the plain black line is its expected distribution under constant size and the dotted line, the expected distribution under growth-decline model. Four indices are specified, mismatch distribution raggedness index r, Fu’s Fs and Ramos-Onsins & Rozas's R_2_ and Tajima’s D. Values noted with in bold are statistically significant * p<0.05, ** P<0.01, *** p<0.001.

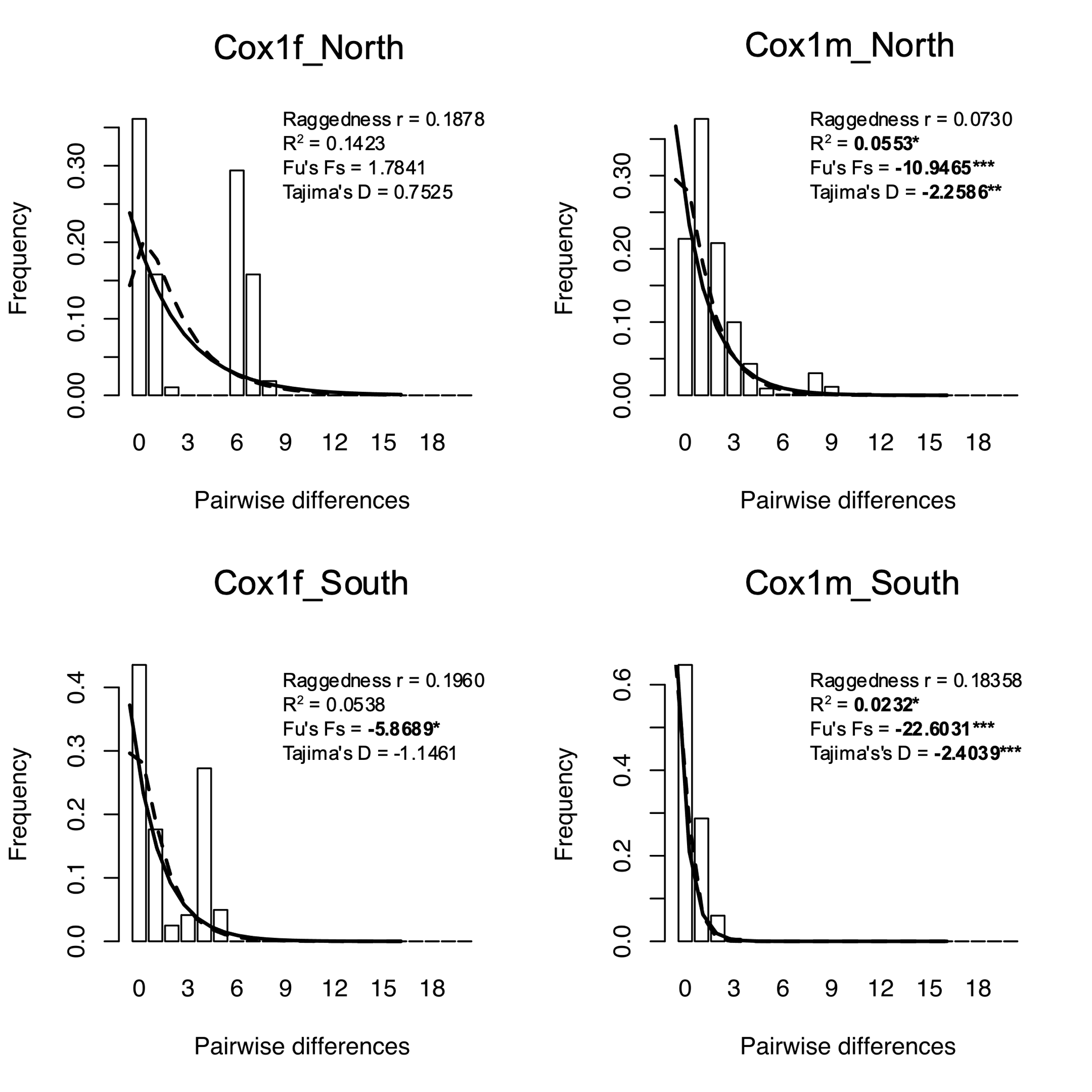
